# Crop and Semi-Natural Habitat Configuration affects Diversity and Abundance of Native Bees (Hymenoptera: Anthophila) in a Large-Scale Cotton Agroecosystem

**DOI:** 10.1101/2020.11.11.377911

**Authors:** Isaac. L. Esquivel, Katherine A. Parys, Karen W. Wright, Micky D. Eubanks, John D. Oswald, Robert N. Coulson, Michael J. Brewer

## Abstract

The cotton agroecosystem is one of the most intensely managed, economically, and culturally important fiber crops worldwide including in the United States of America (U.S.), China, India, Pakistan, and Brazil. The composition and configuration of crop species and semi-natural habitat can have significant effects on ecosystem services such as pollination. Here we investigate the effect of crop and semi-natural habitat configuration in a large-scale cotton agroecosystem on the diversity and abundance of native bees. Interfaces sampled include cotton grown next to cotton, sorghum or semi-natural habitat. Collections of native bees across interface types revealed 32 species in 13 genera across 3 families. Average species richness ranged between 20.5 and 30.5 with the highest (30.5) at the interface of cotton and semi-natural habitat. The most abundant species was *Melissodes tepaneca* Cresson (> 4,000 individuals, ~75% of bees collected) with a higher number of individuals found in all cotton-crop interfaces compared to the cotton interface with semi-natural habitat or natural habitat alone. It was also found that interface type had a significant effect on the native bee communities. Communities of native bees in the cotton-crop interfaces tended to be more consistent in the abundance of species and number of species at each sampling site. While cotton grown next to semi-natural habitat had higher species richness, the number of bees collected varied. These data suggest that native bee communities persist in large-scale cotton agroecosystems and some species may thrive even when cotton-crop interfaces are dominant compared with semi-natural habitat. These data have native bee conservation implications that may improve potential pollination benefits to cotton production.

## Introduction

Anthropogenic intensification has resulted in a simplification of agricultural landscapes or agroecosystems, leaving small fragments of natural habitats among a few dominant crop species. This modification to the landscape is relevant to the agroecosystems of the United States of America (U.S.), Brazil, and other regions where large-scale cotton production occurs. Even in simplified agroecosystems, the composition and configuration of crop species and semi-natural habitat can have significant effects on ecosystem services such as natural pest control and pollination [1]. Semi-natural habitats provide essential resources such as pollen, nectar, alternative hosts, and over-wintering sites for natural enemies and pollinators [1]. These can bolster the ecosystem services they provide which are important for pest management and crop production [1, 2].

Achieving efficient and productive agricultural land use while conserving biodiversity is an important challenge to agricultural sustainability. These issues can be linked. Native bee diversity is relevant to the decline of pollinators in the U.S. This decline may have been affected by large-scale plantings and insecticide use in field crops, such as cotton in the U.S. and other countries [2]. The diversity and abundance of native bee pollinators are important in providing pollination services to a diverse array of crops, many of which receive well understood pollination services from managed bees (including, honey bees, bumble bees, and select commercially managed native bees) or less understood pollination benefits from native bees, which are prevalent in cotton [3,4].

The cotton agroecosystem is one of the most intensely managed, economically, and culturally important fiber crops worldwide. Together, the U.S., China, India, Pakistan, and Brazil comprise the top 5 cotton producers in the world. In the U.S., more than 4.5 million ha of cotton were cultivated in 2017, all of which were planted in the southern U.S. Cotton Belt, spanning from Virginia to California [5]. Texas produces roughly 45% of the U.S. cotton, including where this study is located [5]. Under agricultural intensification, as seen in our model cotton agroecosystem, field sizes commonly exceed 240 hectares.

Pests of cotton, including heliothine pests (Lepidoptera: Noctuidae) and the cotton boll weevil (*Anthonomus grandis* Bohemen, Coleoptera: Curculionidae) were historically controlled using intensive applications of insecticide. With the introduction and development of Bt cotton for control of heliothine pests and area-wide boll weevil eradication efforts, there has been a substantial reduction in foliar insecticide applications for cotton pests in the both the U.S and other cotton growing regions worldwide [6]. This overall reduction of foliar insecticide sprays could be beneficial to local bee communities. Several species of non-*Apis* bees have been observed visiting and nesting in cotton fields frequently [7], and the potential contributions of cotton to aid bee conservation are high [8]. Cotton is perennial plant in the family Malvaceae, which is managed as an annual crop for fiber production. The plants have large flowers that produce large quantities of pollen and nectar that offer both pollen and nectar as resources for many insects, including bees. The availability of mass flowering crops such as cotton across agricultural landscapes often has a positive impact on the density of generalist, native bee species and possibly biodiversity [8].

Yet, information on the pollinator status of native bees and potential benefits to cotton is limited and outdated. Much of the literature on bees and their activity in U.S. cotton is roughly 30 years old, and much of the work was completed prior to the substantial reduction in the use of pesticides in cotton production. Much of this work was done in Arizona and the adjoining Texas Panhandle to identify potential bee species that could be developed and managed for economically feasible hybrid cottonseed production with special attention to the non-native and intensively managed *Apis mellifera* L. [9]. Although cotton is generally considered self-pollinating, previous studies suggest it does benefit from cross-pollination [10]. Cusser et al. [10] documented increased seed cotton weights from bolls produced from flowers that were pollinated by hand compared to bolls from flowers that were self-crossed. Pollen produced by cotton plants is too heavy to move between flowers without transport by insects [11].

Large-scale agricultural production and biodiversity conservation have been traditionally viewed as incompatible [5]. Concepts of a biodiversity-ecosystem service relationship within the context of large-scale agricultural systems can benefit from considering the influence of landscape structure and composition given an individual insect species resource needs and foraging capabilities [12]. Landscape structure may affect diversity of pollinator communities in cotton, including native bees, beetles, and syrphid flies. Higher diversity was found in regions that had a higher abundance of semi-natural habitats intermixed within areas used primarily for cotton production [11].

However, cotton producers growing cotton for fiber do not currently utilize managed pollinators (e.g., *Apis*, *bombus*) to aid production nor use agricultural practices that promote the visitation of wild pollinator communities. Further, corbiculate bees (bees with pollen baskets, primarily *Apis* and *Bombus* in the U.S.) cannot effectively collect pollen from plants in the family Malvaceae and often frequently observed visiting cotton flowers for nectar resources [13]. This loss of effectiveness is often attributed with the length of the spines on cotton pollen which physically interfere with the pollen aggregating process used by honey bees and bumble bees [14]. If cotton is to benefit from pollinators, then native, unmanaged, bees such as *Melissodes tepaneca* (Cresson) and native bee diversity are likely to play a major role [15].

We hypothesize that the configuration of agricultural fields and natural gulf prairie habitat of an extant large-scale cotton agroecosystem affects biodiversity and abundance of native bees. The objective of this study is to investigate the diversity and abundance of native bee pollinators in a model large-scale cotton agroecosystem with consideration of landscape configuration of crops and semi-natural habitat at the local scale. The potential of native bee pollinators benefiting from cotton is considered using the results of this study and placed in context of potential native bee pollination benefits to cotton production.

## Materials and Methods

### Study System

The study area is approximately 12,000 hectares of a large commercial farming operation managed by one private entity that provided continuous access and agronomic management records. This commercial farming operation was located within the south Texas coastal cotton-growing region. The main crops consisted of an annual rotation of upland cotton varieties and sorghum at an approximately 1:1 ratio and were grown following standard agronomic practices for the Texas coastal region [16]. The study area was juxtaposed on the natural gulf prairie habitat consisting of shrubland and a network of rivers, streams, and creeks that drain into the Gulf of Mexico. Field sizes and shapes ranged from 200 to 600 hectares and varied from high curvilinearity with large edge-to-area ratios to simple polygons with low edge-to-area ratios. This allowed for bee collections to be taken at five different cotton-crop or cotton-semi natural habitat interfaces. Bee collections occurred in 2017 and 2018, with specific sample site locations changing between the years. Three interfaces were considered in 2017: a cotton field grown next to another cotton field (designated as cotton-cotton, CC in graphics), cotton grown next to sorghum (designated as cotton-sorghum, CS in graphics), and cotton grown next to semi-natural habitat (designated as cotton-natural habitat, CN in graphics). In 2018, in addition to these three interfaces, two more interface treatments were considered. These were cotton grown next to another cotton field greater than 1 km away from semi-natural habitat (designated as cotton-cotton-far, CCF in graphics) and semi-natural habitat only, not next to cotton (i.e. at least 200m from a cotton field edge, designated as natural habitat, NH in graphics).

### Bee Collection and Processing

Bee bowls (i.e., modified pan traps) were used for collecting native bees. Bee bowls are the most cost-effective and simplest method to monitor bees in agricultural systems [17,18]. These consisted of three ~100 ml (3.25 oz) Solo cups, painted either flat white, fluorescent blue, or fluorescent yellow. The three-bee bowls were individually fastened to shelving brackets held with industrial strength Velcro and attached to T-posts staked into the ground. The bee bowls were positioned at the canopy level. In 2017, three bee bowl units per three selected crop interfaces (Cotton-Cotton, Cotton-Sorghum, Cotton-Semi Natural) were sampled, totaling 27 individual bee bowls for each sampling event. In 2018, five bee bowl units were placed at all interfaces (Cotton-Cotton, Cotton-Sorghum, Cotton-Semi Natural, Cotton-Cotton Far, Natural-Habitat), totaling 75 individual bee bowls per sampling event. Trap units were set out at first bloom and sampled weekly for a period of four weeks for each year. Each week, bee bowl units were randomly placed along the specified interfaces in the morning and collected 24hrs later or soon thereafter when weather and road conditions allowed.

Once bees were collected, specimens were temporarily stored in 70% ethanol, then pinned and labeled following curatorial best practices. Bees were processed by sorting specimens to morphotypes, and later identified to genus using general keys [19,20]. Following is a list of genera and corresponding primary literature used for identifications to species: *Agapostemon* [21], *Anthophora* [22], *Augochlora* [23,24], *Augochlorella*, Ceratina [25,26], *Diadasia* [27], *Halictus* [28], *Lasioglossum* [29,30], *Megachile* [28,31], *Melissodes* and *Svastra* [32–34], *Nomia* [35], and *Xylocopa* [36]. Specimens in the genera *Ceratina* and *Lasioglossum* (*Dialictus*) were not identified to species and left with morphospecies designations due to taxonomic uncertainty within the region.

### Analyses

To characterize the overall diversity of the native bee community within the cotton agroecosystem, data from both 2017 and 2018 were used. Since we sampled three cotton-crop and cotton-semi natural habitat interfaces in 2017 and five in 2018, years were kept separate for further analysis. Community analysis were conducted using R version 3.6.3 “Holding the Windsock” using packages Vegan, and ggplot2 [37–39]. Data ordination was conducted using non-metric multidimensional scaling (nMDS) of Bray-Curtis dissimilarities using Vegan and graphed in ggplot2 to visualize sample and species relationships in a low-dimensional space. Briefly, nMDS arranges points to maximize the rank-order correlation between real-world distances (Bray-Curtis) of species data between sites. It plots the pair-wise dissimilarity between objects in ordination space. Objects (in this case, sites) that are ordinated closer to one another are more similar in species composition and abundance than those further apart. An analysis of similarity test (ANOSIM) using the Bray-Curtis similarity matrix obtained from 999 permutations was used to test differences between the native bee community at different interfaces using the function ‘anosim’ in the R package Vegan [37, 38]. The ANOSIM analysis produces a test statistic ‘R’ that compares the mean of ranked dissimilarities between groups to the mean of ranked dissimilarities within groups. A p-value of the R statistic is determined by multiple permutations of the group membership to obtain the null distribution of the R statistic. Comparing the position of the observed R-value to the null distribution allows an assessment of the statistical significance of R [40]. R values range from –1 to 1 with values closer to 1 indicate strong dissimilarity between groups whereas values close to –1, indicate similarity or more dissimilarity within the groups.

In order to evaluate the adequacy of our bee bowls in capturing the native bee fauna in the cotton agroecosystem, species accumulation curves were produced using the function ‘specaccum’ in the R package BiodiversityR using the expected ‘Coleman’ richness [41,42]. To evaluate species richness, we used a ‘chao1’ estimator to evaluate species richness within sites in the R package ‘Vegan’ [38]. Using standard analysis of variance models, we then analyzed the overall abundance of all native bees as well as the most common native bees across three (2017) and five (2018) interface treatments. When a significant effect was found, we further compared abundances across interfaces with Tukey’s HSD means comparisons at the α = 0.05 significance level.

### Results

In 2017, a total of 897 native bees (excluding 21 *A. mellifera*) were collected using bee bowls representing a total of 28 species. Inspection of species accumulation curves for the 2017 native bee community increased at a high rate and began to level-off as more sites and sampling events were added, although a plateau was not reached (Fig. 1). This led to an increased sampling effort in 2018, including the sampling of two additional interfaces. In 2018, a total of 4,666 native bees (excluding 26 *A. mellifera*) were collected, representing 32 species. The combined 2017 and 2018 species accumulation curve increased exponentially, reaching a plateau indicating that the sampling effort was effective in capturing the native bee species in the area using the bee bowls (Fig. 1). Across both years, a total of 5,563 specimens were collected, representing 32 species in 13 genera and three families (Table 1). Average species richness across sites per interface type ranged from 20.5 to 30.5 combined across both years and was relatively similar across all interfaces (Fig. 2). The two most abundant taxa were *Melissodes* (4,233 individuals) consisting of 2 species, and the subgenus *Lasioglossum* (*Dialictus*) (989 individuals) consisting of 12 species/morphospecies across both years.

**Table 1.**
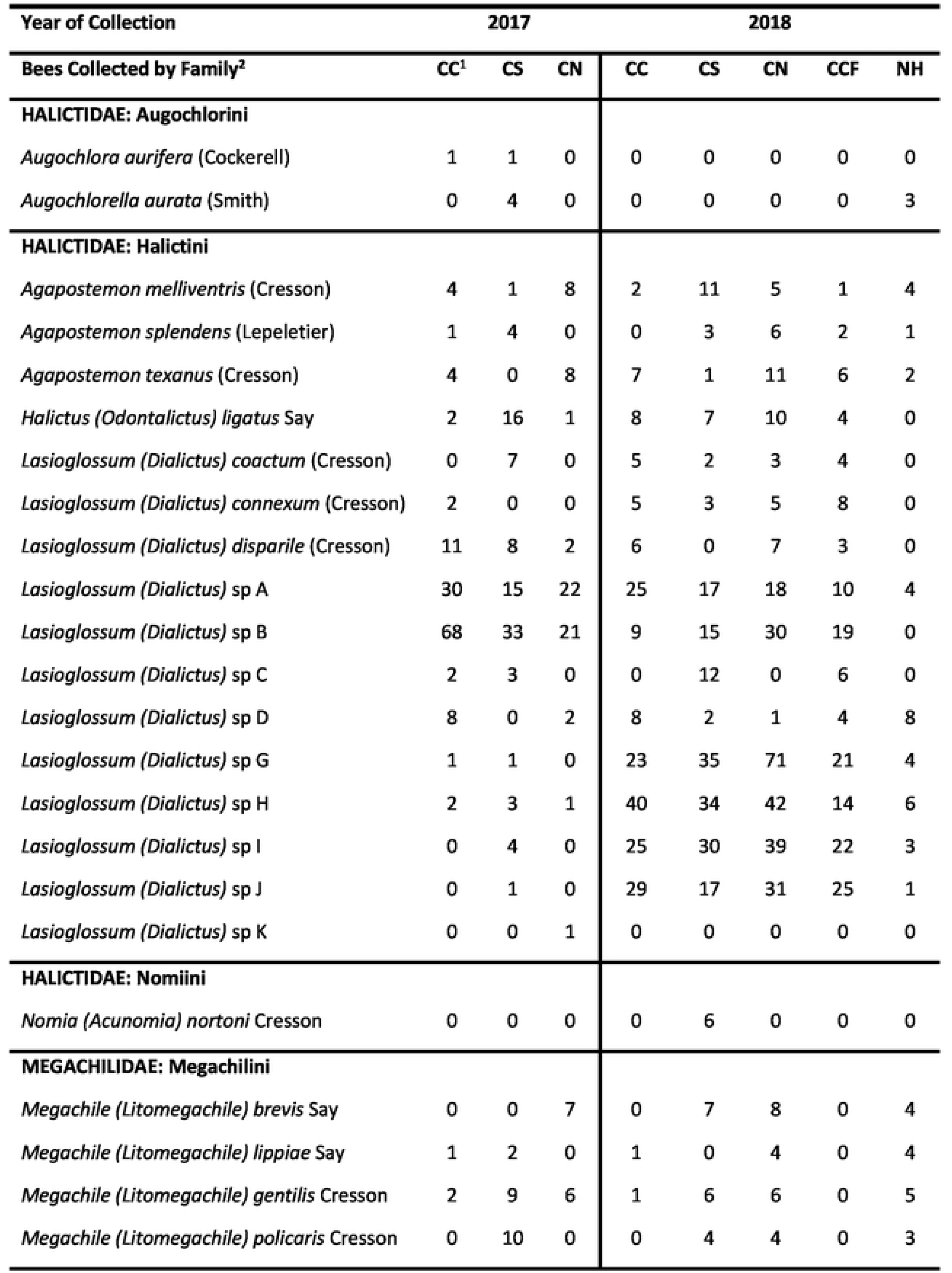
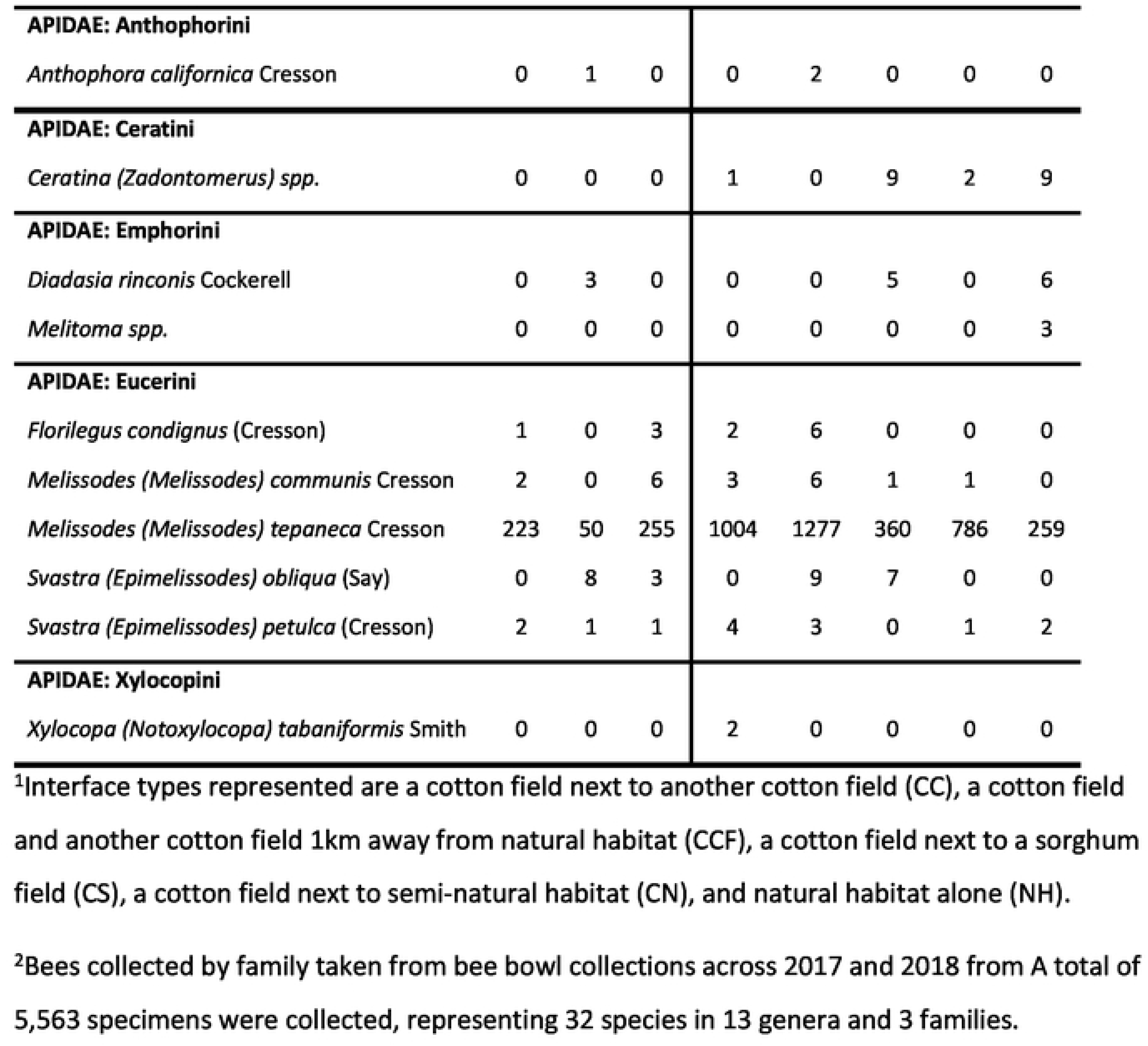
The number of native bees collected at different cottoncrop and cotton semi natural interfaces in a large scale cotton agroecosystem.

**Figure 1.**
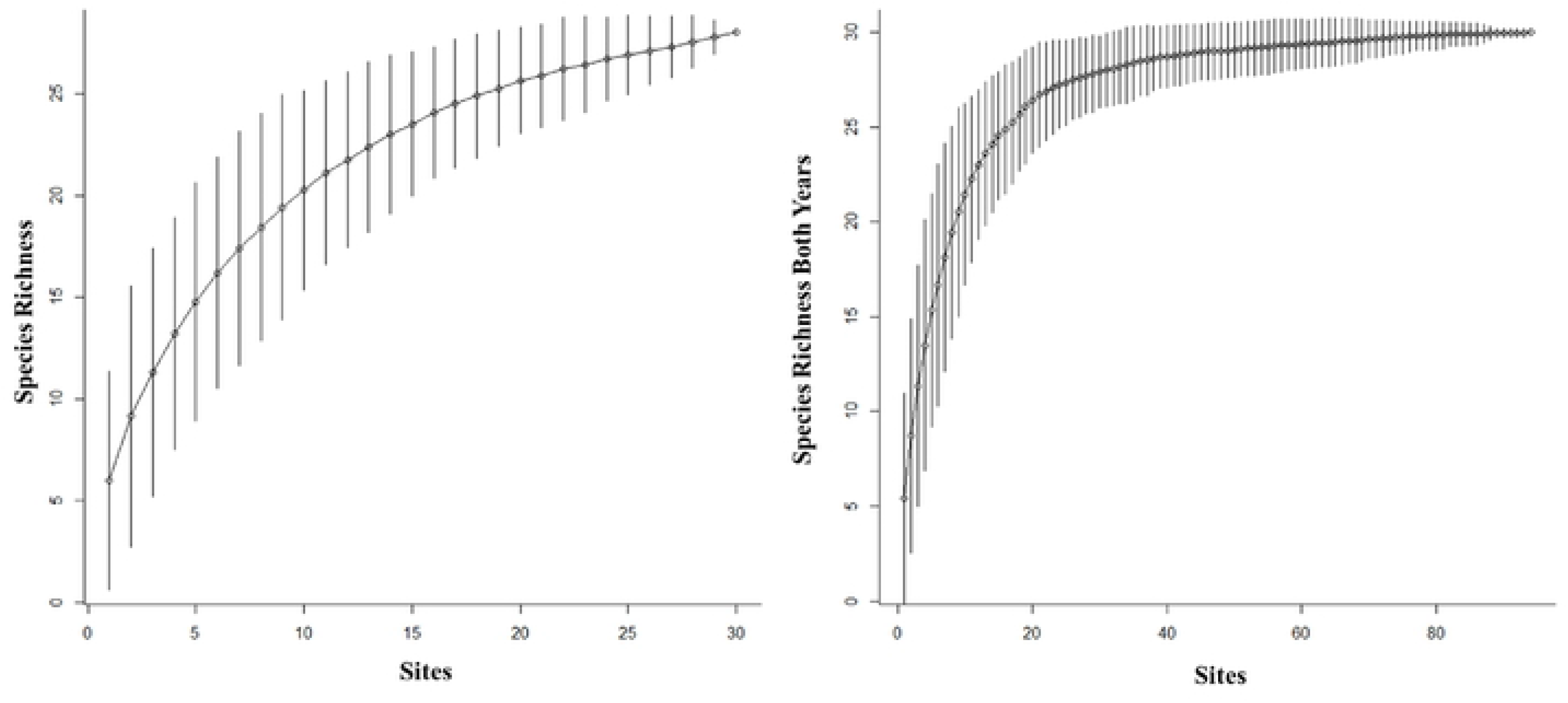
Species accumulation curves from bee bowl collections conducted in 2017 (left) and 2018 (right) across crop and semi-natural habitat interfaces in a large-scale cotton agroecosystem.

**Figure 2.**
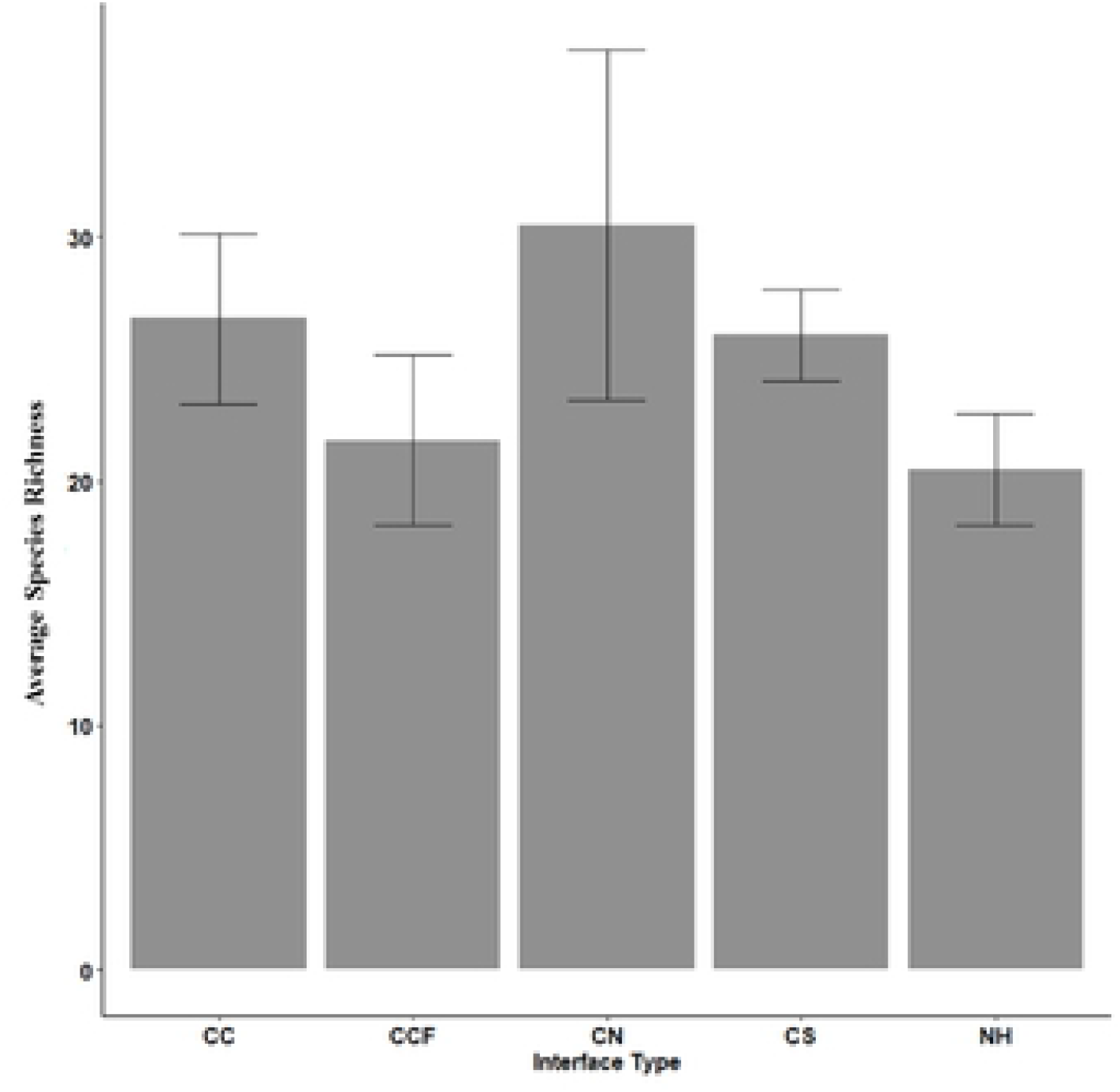
The average species richness calculated from bee-bowl collections in 2017 and 2018 across interfaces from bee-bowl collections in a large-scale cotton agroecosystem. Interface types were sites between a cotton field and another cotton field (CC), a sorghum field (CS), semi-natural habitat (CN), another cotton field 1km away from semi-natural habitat (CCF) and semi-natural habitat alone (NH).

The most common species of the genus *Melissodes* was *Melissodes tepaneca* Cresson, with 528 individuals collected in 2017 and 3686 individuals in 2018. Significant differences in the abundance of *M. tepaneca* were detected across three interfaces in 2017 (*F*= 11.4; df = 2, 27; *P* = 0.0003) and five interface types in 2018 (including semi-natural habitat) (*F*= 5.55; df = 4, 89; *P* < 0.0005) (Fig 3). In 2017, the interface of cotton-semi-natural habitat had the lowest abundance of *M. tepaneca* on average compared to the interfaces of a cotton-cotton, and cotton-sorghum (Fig 3). In 2018, more *M. tepaneca* were collected on average at the interface of a cotton-sorghum compared to the interface of cotton-semi-natural habitat or natural habitat alone. Abundance of *M. tepaneca* in the remaining two interfaces were intermediate in abundance, including the cotton-cotton-far interface that was 1km from natural habitat (Fig. 3).

**Figure 3.**
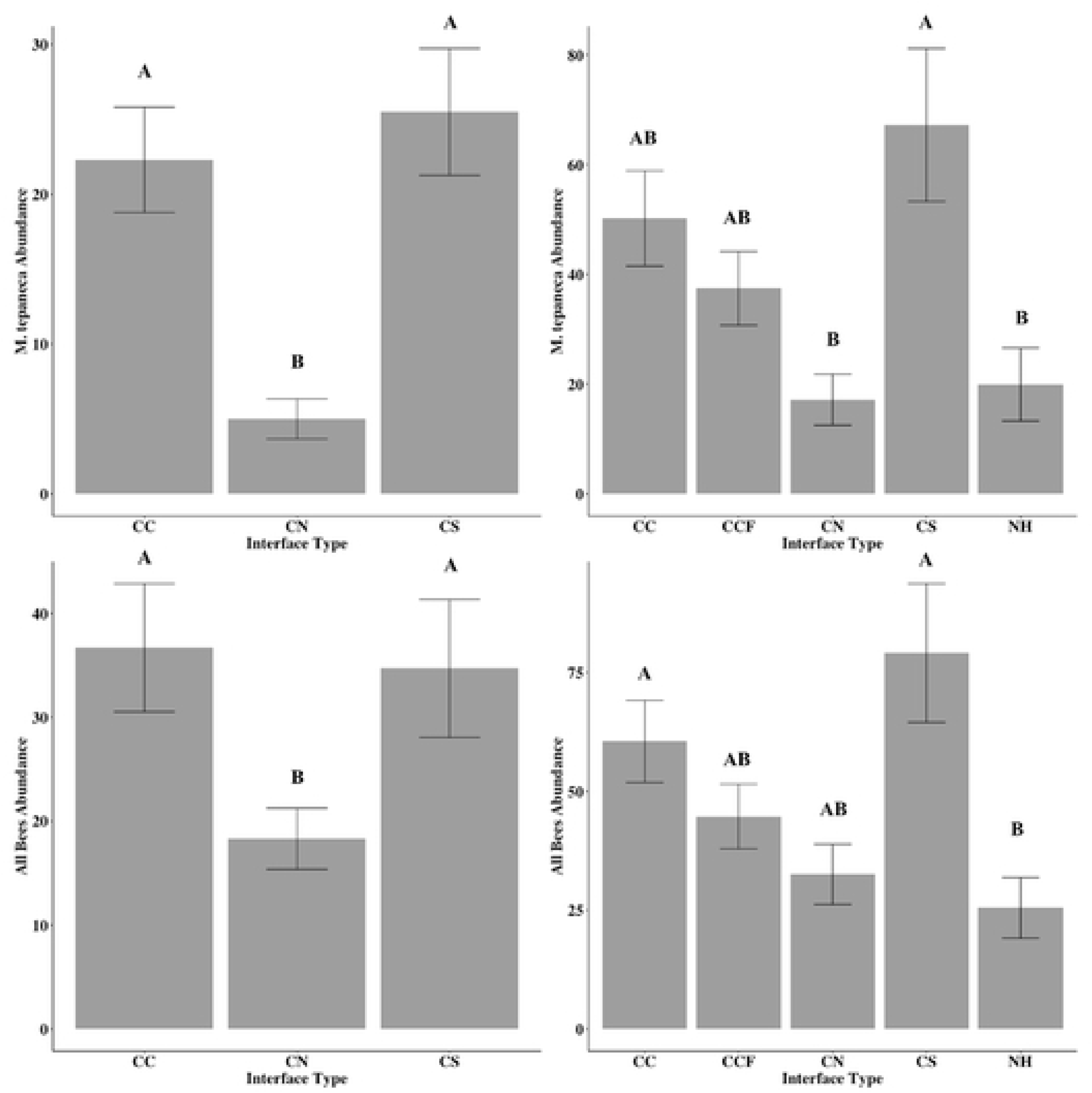
The average abundance and standard error of *Melissodes tepaneca* across crop and semi-natural interfaces taken from bee-bowl collection conducted in 2017 (top left) and 2018 (top right) and abundance of all bee species in 2017 (bottom left) and 2018 (bottom right) in a large-scale cotton agroecosystem. The interface types were sites between a cotton field and another cotton field (CC), a sorghum field (CS), semi-natural habitat (CN), another cotton field 1km away from semi-natural habitat (CCF) and semi-natural habitat alone (NH). Abundances across interface types were compared using Tukey’s HSD means procedure with letters differing across means indicated significant differences.

Looking at the total amount of native bees collected in both 2017 (*F*= 3.37; df = 2, 27; *P* < 0.044) and 2018 (*F*= 5.13; df = 4, 89; *P* = 0.0009), the results were the same as when analyzing the *M. tepaneca* data only. This similarity at least partially reflects the high frequency and evenness of occurrence of *M. tepaneca* in the collections of all interface types. In 2017, more native bees were collected on average at the interface of cotton-sorghum and cotton-cotton compared to the interface of a cotton field-semi-natural habitat (Fig. 3). In 2018, more native bees on average were collected at the interface of a cotton-sorghum, and cotton-cotton, compared to cotton-natural habitat alone (Fig. 3) The interfaces of a cotton-cotton-far and cotton-semi-natural habitat were intermediate in abundances (Fig. 3).

Interface type had a significant effect on the abundance of 12 *Lasioglossum* (*Dialictus*) species in both 2017 (*F*= 3.68; df = 2, 27; *P* = 0.0403) and 2018 (*F*= 2.99; df = 4, 89; *P* = 0.02361) (Fig. 4). In 2017, more *Lassioglosum* (*Dialictus*) spp. were collected on average at the interface of cotton-cotton compared to the interface of cotton-sorghum. The interface of cotton-semi-natural habitat had an intermediate amount of *Dialictus spp.* (Fig. 3). In 2018, more *Lassioglosum* (*Dialictus)* spp. were caught on average the cotton and semi-natural habitat interface compared to the natural habitat alone. The other interfaces had an intermediate number of *Dialictus spp*. collected (Fig. 4).

**Figure 4.**
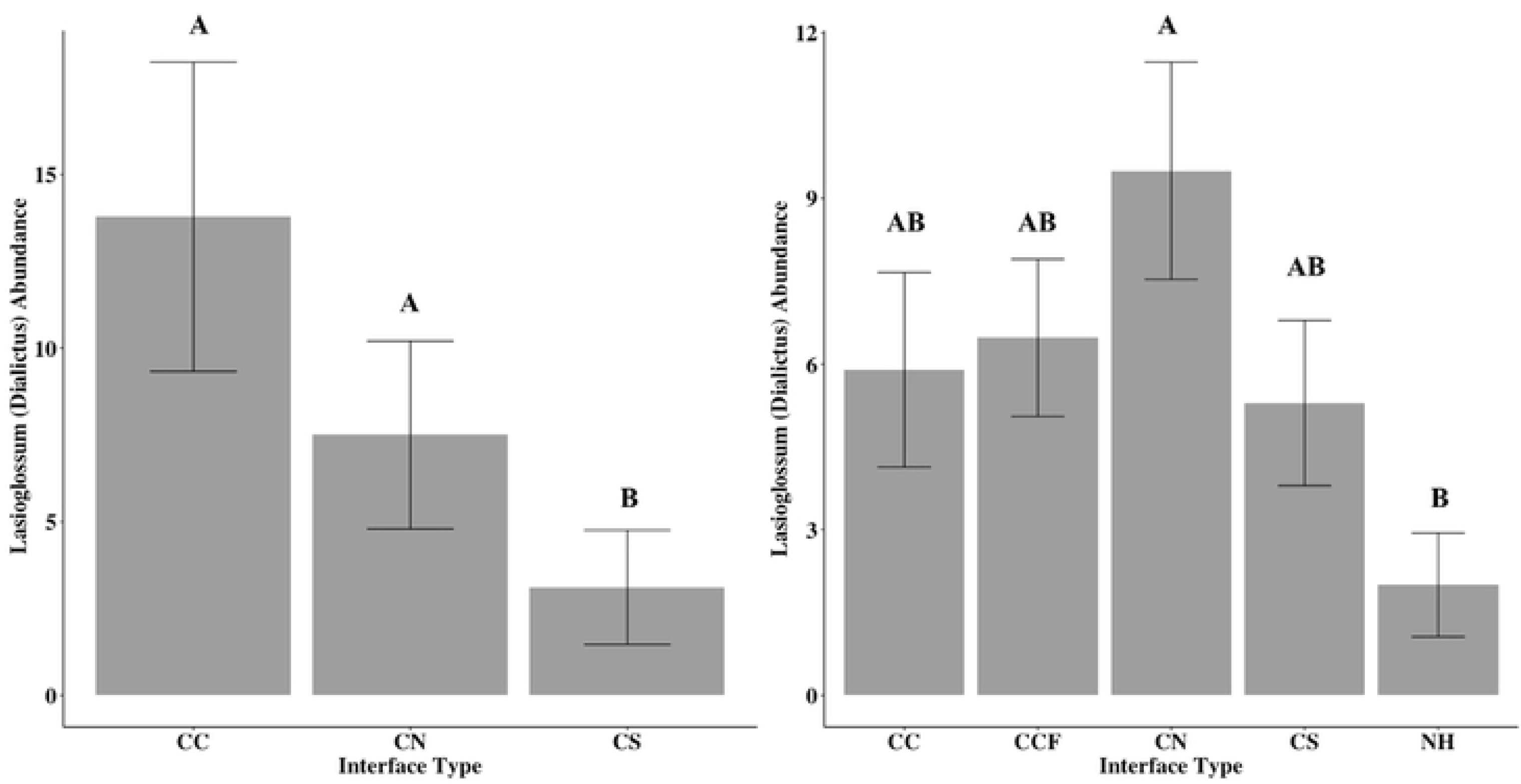
Average abundance and standard error *of Dialictus (Lassioglossum) spp.* across crop and semi-natural interfaces taken from bee-bowl collection conducted in 2017 and 2018 in a large-scale cotton agroecosystem. The interface types were sites between a cotton field and another cotton field (CC), a sorghum field (CS), semi-natural habitat (CN), another cotton field 1km away from semi-natural habitat (CCF) and semi-natural habitat alone (NH). Abundances across interface types were compared using Tukey’s HSD means procedure with letters differing across means indicated significant differences.

In 2017, three groups of native bee communities of the three interface types appeared distinctive, showing well-formed groups on the nMDS plot (Fig. 5). The reasonably low-stress level (0.0768) indicated a fair representation of multidimensional space. The ANOSIM analysis indicated moderate differences (R= 0.2914) and was overall significant (*P* = 0.001), indicating that the communities between the sampling sites across the cotton-cotton, cotton-sorghum, and cotton-semi-natural habitat interfaces were more different from one another, while the sampling sites within each interface type were more similar to each other. The width of the ellipses suggests that sites in the cotton-natural habitat interface had a higher variance in the similarity between sites. In contrast, the other two interfaces were more consistent in the abundance of each species found. In 2018, the nMDS ordination indicated groupings of communities between the five interface types with a reasonably low-stress score (0.0944). However, the confidence ellipses with an ANOSIM test statistic (R= 0.1511) showed some overlap between the interface groupings with at least some differences in the groups detected (*P* = 0.001) (Fig. 5). This is reflected in the position and the size of the ellipse groupings in the nMDS plot (Fig. 5). For example, the natural habitat ellipse appears to overlap all the other communities of the cotton-crop and cotton-natural interfaces. This suggests that although most species are found in the natural habitat alone, the abundance of individuals caught between sampling sites across the samples was more variable. Whereas the ellipses at the interface of cotton-sorghum and cotton-cotton were rather tight, suggesting that there was consistency in the abundance and number of bee species collected between the sampling sites at each sampling event.

**Figure 5.**
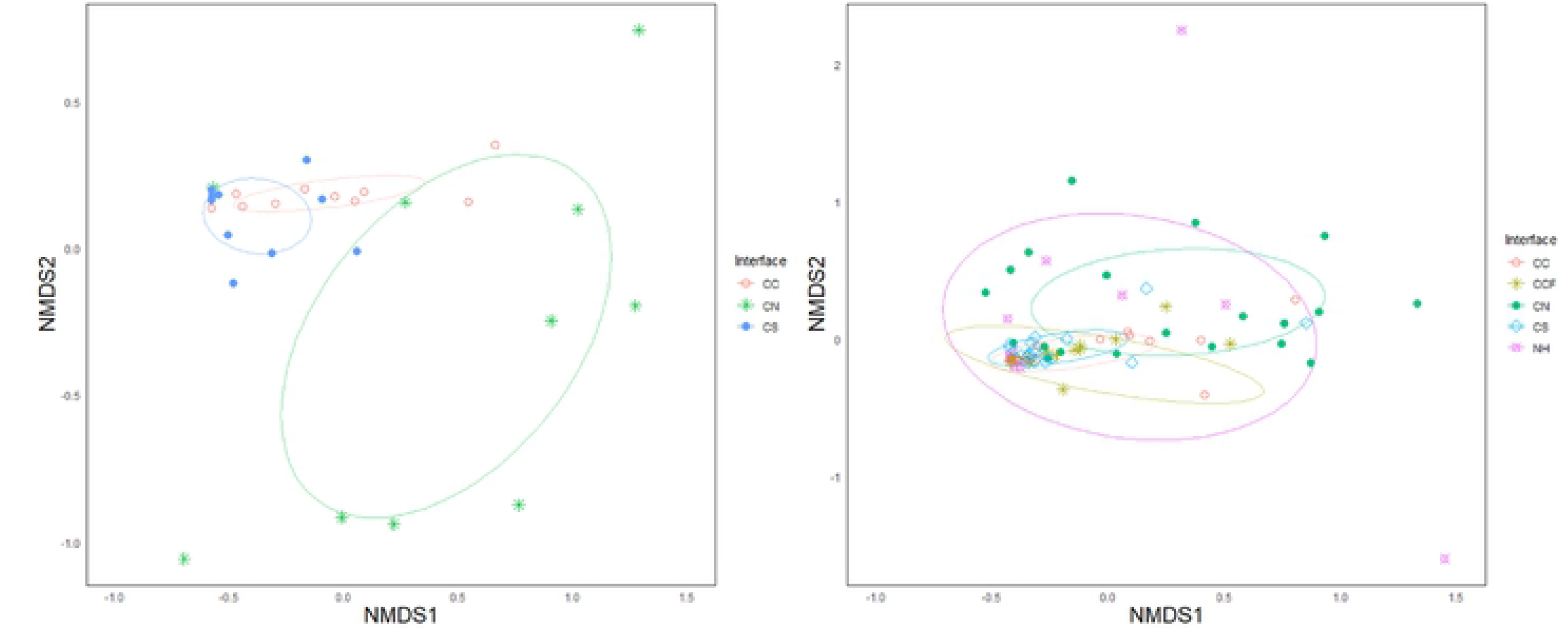
A Non-metric multidimensional scaling (NMDS) of Bray-Curtis similarity data of native bee communities in 2017 (left) and 2018 (right) across crop and semi-natural interfaces in a large-scale cotton agroecosystem. Each point is representative of a site placed according to their similarities with other sites, and ellipses representing the variance of the abundance of species within those sites. Widths of ellipses indicating the bee community similarity within sites at a given interface are greater than the similarities between the sites at different interface

## Discussion

In the large-scale cotton agroecosystem, a total of 33 species (including *A. mellifera*) were collected, with a total of 5,563 native bees collected across 2017 and 2018. The majority of these bees were composed of a single species, *Melissodes tepaneca*, representing 75% of the individuals collected. Bee species found in this study have diverse nesting preferences. While bees in the genus *Megachile* are strictly solitary, some species of *Lassioglosum* (*Dialictus*) are semi-social, and others in genera like *Melissodes* and *Nomia* are solitary, yet gregarious [19]. Most of these species are generalist pollinators except for *Florilegus condignus* (Cresson), a known specialist on *Pondenteria spp*. amongst others, and *Diadasia rinconis* (Cockerell), which is eclectically oligolectic and a specialist of *Opuntia spp.* and *Spharalcea spp*. Interestingly, plants in the genus *Spharalcea* are in the same family as cotton, Malvaceae, yet *D. rinconis* is quite rare in our system.

These data are consistent with both past and current literature in which species in the genus *Melissodes* are found within cotton fields in various parts of U.S cotton-growing regions. For example, in experimental observations of bees visiting Georgia cotton flowers by Allard (1911), roughly 80% of over 2,000 visual identifications were of a single species, *Melissodes bimaculatus* (Lepeltier). *Melissodes thelypodii* (Cockerell) was also abundant in cotton of the Texas High Plains [9]. In the same study, one species in the genus *Agapostemon, A. angelicus* (Cockerell), was found in high abundance [9]. In our study, we found three species of *Agapostemon* at detectable rates but low abundance. More recently in Texas, using hand collection methods, 37 species of bees were found in small-scale cotton systems with most field sizes below 200 ha. At this small scale, 21% out of 800 total specimens collected were *M. tepaneca* [10]. The genus *Melissodes* also appears to be important in cotton outside of the U.S. In Brazil, *M. nigroaena* (Smith) has been identified as an important pollinator in small-scale cotton agroecosystems systems [43,44].

Unlike some of the studies previously mentioned where native bee abundances and diversity were measured at smaller scale cotton fields (< 50 hectares) with more diverse landscapes, our study took place on large-scale commercial cotton fields where average field sizes were approximately 240 hectares. Semi-natural habitats were relatively sparse and primarily associated with a natural and partially augmented (i.e. drainage ditches along cultivated fields) water system flowing into the Gulf of Mexico. Shrubland and pasture where cattle graze were also found in large continuous tracks of land nearby. Agricultural intensification in our study area also included tillage for planting and weed control and insecticide applications for various cotton and sorghum pests, although insecticide use was substantially reduced from previous decades [6]. Despite the dominance of the crop-crop interfaces and other aspects of agricultural intensification, the bee communities were similar in terms of species richness at different interfaces of this highly structured large-scale cotton agroecosystem, suggesting benefits or at least no antagonism of crops intermixed and dominating the original natural gulf prairie habitat. Compared to other studies, where increasing native bee diversity and richness in more diverse landscapes was seen, our findings were rather unexpected given the higher diversity and abundance within agricultural interfaces [10,43,44]. Further, samples taken from natural habitat alone (NH) appeared to have a lower estimated species richness than all cotton-crop or cotton-natural habitat interfaces, supporting that flowering cotton and possibly other resources provided by sorghum benefited at least *M. tepaneca* in this agroecosystem that was designed from a cotton production perspective.

Although species composition was similar across interfaces and semi-natural habitat, more native bees overall were collected within cotton-crop interfaces. Specifically, *M. tepaneca* appears to have an affinity for crops as it was collected in higher amounts at interfaces of cotton-crop compared to the interface of cotton-semi-natural habitat, or semi-natural habitat alone. They were most abundant at the interface of a cotton-sorghum, in both years. Abundance of *M. tepaneca* was similar at the among the cotton-cotton interfaces and the cotton-cotton-far interfaces. The average foraging distance from nesting site to food site for most solitary bees is between 150-600m depending on body size [45]. In a cotton operation where fields are on average 500-1000m wide, this suggests that at least for *M. tepaneca* nesting occurs within the area of agricultural intensification with few remnants of semi-natural habitat. It has been documented the *M. tepaneca* does nest within cotton rows in Arizona cotton fields [46] which may be representative of *M. tepaneca* in Texas cotton fields. Several species within the genus *Melissodes* have been observed to nest in aggregations of up to 200 individual nests in semi-sandy soils, both within patches devoid of vegetation and partially covered ground with leaflitter or grass [47,48].

On the other hand, *Lasioglossum* (*Dialictus*) spp. were more abundant at the cotton-semi-natural habitat interface. Further, less *Lasioglossum* (*Dialictus*) were found in the natural habitat alone compared to the interface of cotton-semi-natural habitat. This suggests that some of the species in this group have an affinity to cotton possibly utilizing a resource in cotton when in cultivation. This group is semi-social and highly diverse in abundance and life history. Species range specialists to generalists, and display a variety of life histories, inclusive of parasitism, and may be uncommon to abundant [30]. More detailed research on foraging and nesting behavior of these species in agroecosystems is warranted to explore the mechanisms that may be driving our observation that native bee community within a large-scale cotton agroecosystem persists, and some species may thrive even when crop-crop ecotone interfaces dominate semi-natural habitat.

We have shown that a native bee community is persistent amongst various interfaces of crops and semi-natural habitat across large-scale cotton agroecosystems and shows similarities to other cropping systems, typically seen at smaller scales [49]. A few of the abundant generalist species found here, including *Melissodes*, *Agapostemon*, *Halictus*, *Lasioglossum (Dialictus)*, and *Nomia*, are also common in other cropping systems across North America, including corn, soybeans, hemp, and alfalfa [17, 18, 50]. This suggests that selected generalist bee species are quite resilient and adaptable to changes caused by agricultural intensification. In comparison rare or threatened species are seldom seen in agricultural fields. Common crops in large scale agroecosystems are often wind pollinated and therefore scarce in pollinator resources, but mass flowering crops such as cotton are an important exception. Loss of specialist bees in these systems is predicted (and consistent with our observations in cotton), but several generalist bees persist and one, *M. tepaneca* appears to thrive.

Bee conservation efforts tend to focus on native or slightly modified habitats. In the U.S. and the European Union, there are various stewardship programs that support portions of agricultural production temporarily or permanently taken out of production to augment wildlife, preserve diversity, and aid pollinators and other organisms that provide ecosystem services and eco-tourism [51]. As stated by Tscharntke et al. (2005), there has been a recent debate that this type of conservation to augment ecosystem services is of limited value when considering the often neglected influence of landscape context on local processes [52–54]. In this case, processes such as crop configuration of the cotton agroecosystem affect native bee pollinators and may offset negative aspects of agricultural intensification. The result is selective pollinators such as *M. tepaneca* may persist and thrive. Further, intensified land use in agriculture and forestry is, without a doubt, a contributor to global climate change and biodiversity loss. However, Tscharntke et al. [6] state biodiversity conservation focusing on 5% of remaining pristine natural habitats will have little value without a recognition of the contribution of the of all land-use types, including intensely managed agricultural land. In the model system studied here, stewardship of native bees appears to be present in a large-scale cotton agroecosystem at the local field-to-field level of scale that may have implications of biodiversity conservation at greater scales.

In a large meta-analysis of pollination services provided by native bee species, Kleijn et al. [55] found that crop-visiting native bee communities are dominated by a small number of common species that persist under agricultural expansion. They also state that many of these pollinators have the potential to be enhanced by simple conservation measures such as modifying tillage mechanisms to no-till systems. Kleijn et al. [55] state these should be the focus of conservation efforts to bolster pollination services provided to agricultural production. In our study, *M. tepaneca* is clearly the most abundant species in the large-scale cotton agroecosystem. This species may benefit from simple conservation measures as well. However, given its abundance in all the cotton-crop interfaces, including cotton-cotton and cotton-cotton-far, it appears to thrive in the extant large-scale system. The scale and quality of crop set-asides and conservation easments are current considerations in conservation programs [56]. The configution of crops and semi-natural habitat in agroecosytems and key species contributors to ecosystem services shoud be added to the debate on pursing bee conservation.

Specific to the cotton agroecosytem, *M. tepaneca* has a positive effect on cotton production by increasing lint and seed weight when flowers are exposed to *M. tepaneca* [15]. The findings here suggest that cotton may benefit *M. tepaneca* and other native bees both near and relatively far from semi-natural habitat. This complementary benefit to cotton supports consideration of a joint bee conservation and cotton productivity approach in at least this select large-scale cotton agroecosystem that has substantially modified the original natural gulf prairie habitat and that was designed from a cotton production perspective. We pose that this system may represent a win-win for bee conservation and cotton productivity, and such a system may be a valuable learning arena to explore underlying mechanisms that benefit bee conservation and augment ecosystem service in the form of pollination [15]. Additional studies into bee natural history in this system such as nesting habitats and preferences, and foraging preferences of *M. tepaneca,* and other common and abundant bee species will aid in understanding of how these bees persist and possibly thrive in intensified agricultural systems of large-scale cotton production.

## Acknowledgments

We thank the C. Everett Salyer Fellowship for Cotton Research (Provided to I.L.E) and USDA NIFA, Southern I.P.M. Center, Enhancement grant Program (Provided to I.L.E). We also thank Cotton the land managers who allowed us access to their property to conduct this study. Additionally, we thank everyone who helped with field collection and processing of bee specimens, especially Miles Arceneaux, who was critical in specimen processing, and Ashleigh Faris.

## References

1. Tscharntke T, Rand T a, Bianchi FJJ a. The landscape context of trophic interactions: insect spillover across the crop-noncrop interface. Ann Zool Fennici. 2005;42: 421–432.

2. Tscharntke T, Clough Y, Wanger TC, Jackson L, Motzke I, Perfecto I, et al. Global food security, biodiversity conservation and the future of agricultural intensification. Biol Conserv. 2012;151: 53–59. doi:10.1016/j.biocon.2012.01.068

3. Klein AM, Vaissière BE, Cane JH, Steffan-Dewenter I, Cunningham SA, Kremen C, et al. Importance of pollinators in changing landscapes for world crops. Proc R Soc B Biol Sci. 2007;274: 303–313. doi:10.1098/rspb.2006.3721

4. Sardiñas HS, Kremen C. Pollination services from field-scale agricultural diversification may be context-dependent. Agric Ecosyst Environ. 2015;207: 17–25. doi:10.1016/j.agee.2015.03.020

5. National Agricultural Statistics Service: Quick Stats. Available online: http://www.nass.usda.gov/Quick_Stats/Lite/ (accessed on 1 January 2020).

6. Luttrell, R.G.; Teague, T.G.; Brewer, M.J. Cotton insect pest management. In Cotton; Fang, D.D., Percey, R.G., Eds.; American Society of Agronomy, Crop Science Society, Soil Science Society of America: Madison, WI, USA, 2015; pp. 509–546

7. Parys KA, Esquivel IL, Wright KW, Griswold T, Brewer MJ. Native pollinators (Hymenoptera : Anthophila) in cotton grown in the Gulf South, United States. 2020; 1–14.

8. Westphal C, Steffan-Dewenter I, Tscharntke T. Mass flowering crops enhance pollinator densities at a landscape scale. Ecol Lett. 2003;6: 961–965. doi:10.1046/j.1461-0248.2003.00523.x

9. Moffett JO, Stith LEES, Burkhardt CC, Shipman CW. Fluctuation of wild bee and wasp visits to cotton flowers. J Arizona Acad Sci. 1976;11: 64–68.

10. Cusser S, Neff JL, Jha S. Natural land cover drives pollinator abundance and richness, leading to reductions in pollen limitation in cotton agroecosystems. Agric Ecosyst Environ. 2016;226: 33–42. doi:10.1016/j.agee.2016.04.020

11. Free JB. Insect pollination of crops. London, UK: Academic Press; 1993.

12. Peterson G, Allen CR, Holling CS. Ecological Resilience, Biodiversity, and Scale. Ecosystems. 1998;1: 6–18. doi:10.1007/s100219900002

13. Vaissière BE, Vinson SB. Pollen morphology and its effect on pollen collection by honey bees, apis mellifera L. (hymenoptera: Apidae), with special reference to upland cotton, gossypium hirsutum L. (malvaceae). Grana. 1994;33: 128–138. doi:10.1080/00173139409428989

14. Konzmann S, Koethe S, Lunau K. Pollen grain morphology is not exclusively responsible for pollen collectability in bumble bees. Sci Rep. 2019;9: 1–8. doi:10.1038/s41598-019-41262-6

15. Esquivel, I. L., R. N. Coulson, and M. J. Brewer. A native bee, *Melissodes tepaneca* (Hymenoptera: Apidae), benefits cotton production. Insects 2020; 1 (8): https://doi.org/10.3390/insects11080487

16. Morgan, G., 2018. Cotton.tamu.edu. Texas A&M AgriLife Extension, College Station (accessed online at http://cotton.tamu.edu/index.html. June 5, 2018).

17. Gill KA, O’Neal ME. Survey of soybean insect pollinators: community identification and sampling method analysis. Environ Entomol. 2015;44: 488–498. doi:10.1093/ee/nvv001

18. Wheelock MJ, Rey KP, O’Neal ME. Defining the insect pollinator community found in iowa corn and soybean fields: implications for pollinator conservation. Environ Entomol. 2016;45: 1099–1106. doi:10.1093/ee/nvw087

19. Michener CD. The bees of the world. Baltimore, MD: John Hopkins University Press; 2007.

20. Michener CD, MacGinley R, Danfoorth B. The bee genera of North and Central America. 1994.

21. Roberts RB. Bees of northwestern America : *Agapostemon* (Hymenoptera: Halictidae). Agric Exp Stn Oregon State Tech Bull. 1973.

22. Cresson ET. A List of the North American species of the genus Anthophora, with descriptions of new species. Trans Am Entomol Soc. 1868;2: 289–293. doi:10.2307/25076210

23. Sandhouse GA. The American bees of the subgenus *Halictus*. Entomol Am. 1941;21: 23–39.

24. Mitchell, T.B. Bees of the Eastern United States (I). North Carolina Ag. Exp. Sta. Bull. 1960, 141, 1–538

25. Daly H V. Bees of the genus *Ceratina* in america north of mexico. U California Pub Entomol. 1973; 172–175.

26. Rehan SM, Sheffield CS. Morphological and molecular delineation of a new species in the *Ceratina dupla* species-group (Hymenoptera: Apidae: Xylocopinae) of eastern North America. Zootaxa. 2011;50: 35–50. doi:10.11646/zootaxa.2873.1.3

27. Sipes SD, Wolf PG. Phylogenetic Relationships within Diadasia, a Group of Specialist Bees. Mol Phylogenet Evol. 2001;19: 144–156. doi:https://doi.org/10.1006/mpev.2001.0914

28. Mitchell TB. Bees of the eastern United States Vol II. Tech Bull 141. 1960;1: 513–546. Available: http://www.discoverlife.org/mp/20q?search=Osmia+albiventris

29. Gibbs J. Revision of the metallic *Lasioglossum* (*Dialictus*) of eastern North America (Hymenoptera: Halictidae: Halictini). Zootaxa. 2011. doi:10.11646/zootaxa.3073.1.1

30. Gibbs J, Packer L, Dumesh S, Danforth BN. Revision and reclassification of *Lasioglossum* (*Evylaeus*), *L*. (*Hemihalictus*) and *L*. (*Sphecodogastra*) in eastern North America (Hymenoptera: Apoidea: Halictidae). Zootaxa. 2013. doi:10.11646/zootaxa.3672.1.1

31. Sheffield CS, Ratti C, Packer L, Griswold T. Leafcutter and Mason Bees of the genus Megachile Latreille (Hymenoptera: Megachilidae) in Canada and Alaska. Can J Arthropod Identif. 2011;18: 1–107. doi:10.3752/cjai.2011.18

32. LaBerge WE. A revision of the bees of the genus Melissodes in North and Central America. Part I. Univ Kansas Sci Bull. 1956;37: 911–1194.

33. LaBerge WE. A revision of the bees of the genus Melissodes in North and Central America. Part II. Univ Kansas Sci Bull. 1956;38: 533–578.

34. LaBerge WE. A revision of the bees of the genus Melissodes in North and Central America. Part III. Univ Kansas Sci Bull. 1961;42: 283–663.

35. Cockerell TDA. The North American bees of the genus Nomia. 1911. Available: file://catalog.hathitrust.org/Record/100359478

36. Hurd PD. The carpenter bees of california. Bull Calif Insect Surv. 1955;4: 35–72. doi:10.1017/CBO9781107415324.004

37. Wickham H. ggplot2: Elegant Graphics for Data Analysis. New York: Springer-Verlag; 2016. Available: https://ggplot2.tidyverse.org

38. Oksanen, J.; Blanchet, F.G.; Friendly, M.; Kindt, R.; Legendre, P.; McGlinn, D.; Minchin, P.R.; O’Hara, R.B.; Simpson, G.L.; Solymos, P.; et al. VEGAN: Community Ecology Package. R Package Version 2, 5–6. 2019. Available online: https://CRAN.R-project.org/package=vegan (accessed on 14 May 2020).

39. R Core Team. R: A language and environment for statistical computing. Vienna, Austria: R Foundation for Statistical Computing; 2020. Available: https://www.r-project.org/

40. Clarke KR. Non‐parametric multivariate analyses of changes in community structure. Aust J Ecol. 1993;18: 117–143. doi:10.1111/j.1442-9993.1993.tb00438.x

41. Gotelli NJ, Colwell RK. Quantifyinf Biodiversity: Procedures and pitfalls in the measurement and comparison of species richness. Ecol Lett. 2001;4: 379–391. doi:10.1046/j.1461-0248.2001.00230.x

42. Kindt, R. Coe R. Tree diversity analysis. A manual and software for common statistical methods for ecological and biodiversity studies. Nairobi (Kenya): World Agroforestry Centre (ICRAF); 2005. Available: http://www.worldagroforestry.org/output/tree-diversity-analysis

43. Grando C, Amon ND, Clough SJ, Guo N, Wei W, Azevedo P, et al. Two colors, one species: The case of *Melissodes nigroaenea* (Apidae: Eucerini), an important pollinator of cotton fields in Brazil. Sociobiology. 2018;65: 645–653. doi:10.13102/sociobiology.v65i4.3464

44. Cusser S, Grando C, Zucchi MI, López-Uribe MM, Pope NS, Ballare K, et al. Small but critical: semi-natural habitat fragments promote bee abundance in cotton agroecosystems across both Brazil and the United States. Landsc Ecol. 2019;34: 1825–1836. doi:10.1007/s10980-019-00868-x

45. Gathmann A, Tscharntke T. Foraging ranges of solitary bees. J Anim Ecol. 2002;71: 757–764. doi:10.1046/j.1365-2656.2002.00641.x

46. Butler, G.d., Todd, F.E., McGregor, S.E., Werner FG. Melissodes bees in Arizona cotton field. Agric Exp Stn Univ Arizona. 1960;3: 1–11.

47. Clement SL. The Nesting Biology of *Melissodes* (Eumelissodes) *rustica* (Say), with a description of the larva. Published by : Allen Press on behalf of Kansas (Central States) Entomological Society content in a trusted digital archive. 2011; 46: 516–525.

48. Cameron ASA, Whitfield JB, Hulslander CL, Cresko WA, Isenberg SB, King RW. Nesting biology and foraging patterns of the solitary bee *Melissodes rustica* (Hymenoptera : Apidae) in Northwest Arkansas Source : Journal of the Kansas Entomological Society Vol. 69, No. 4, Supplement : Special Publication Number 2 : Proceedings. 2013;69.

49. Hladik ML, Vandever M, Smalling KL. Exposure of native bees foraging in an agricultural landscape to current-use pesticides. Sci Total Environ. 2016;542: 469–477. doi:10.1016/j.scitotenv.2015.10.077

50. O’Brien C, Arathi HS. Bee genera, diversity and abundance in genetically modified canola fields. GM Crop Food. 2018;9: 31–38. doi:10.1080/21645698.2018.1445470

51. Brewer MJ, Goodell PB. Approaches and incentives to implement integrated pest management that addresses regional and environmental issues. Annu Rev Entomol. 2012;57: 41–59. doi:10.1146/annurev-ento-120709-144748

52. Collins WW, Qualset CO. Biodiversity in agroecosystems. CRC Press, Boca Raton, USA; 1999.

53. Bengtsson J, Angelstam P, Elmqvist T, Emanuelsson U, Folke C, Ihse M, et al. Reserves, resilience and dynamic landscapes. AMBIO A J Hum Environ. 2003;32: 389–396. doi:10.1579/0044-7447-32.6.389

54. Kleijn D, Winfree R, Bartomeus I, Carvalheiro LG, Henry M, Isaacs R, et al. Delivery of crop pollination services is an insufficient argument for wild pollinator conservation. Nat Commun. 2015;6. doi:10.1038/ncomms8414

55. Kleijn D, Winfree R, Bartomeus I, Carvalheiro LG, Henry M, Isaacs R, et al. Delivery of crop pollination services is an insufficient argument for wild pollinator conservation. Nat Commun. 2015;6. doi:10.1038/ncomms8414

56. Kleijn D, Baquero RA, Clough Y, Díaz M, De Esteban J, Fernández F, et al. Mixed biodiversity benefits of agri-environment schemes in five European countries. Ecol Lett. 2006;9: 243–254. doi:10.1111/j.1461-0248.2005.00869.x

